# Bacterial microcompartments for isethionate desulfonation in the taurine-degrading human-gut bacterium *Bilophila wadsworthia*

**DOI:** 10.1101/2021.07.13.451622

**Authors:** Anna G. Burrichter, Stefanie Dörr, Paavo Bergmann, Sebastian Haiß, Anja Keller, Corentin Fournier, Paolo Franchini, Erika Isono, David Schleheck

**Author notes:** Correspondence authorships: Anna Burrichter, and David Schleheck.

## Abstract

**Background:** *Bilophila wadsworthia*, a strictly anaerobic, sulfite-reducing bacterium and common member of the human gut microbiota, has been associated with diseases such as appendicitis and colitis. It is specialized on organosulfonate respiration for energy conservation, i.e., utilization of dietary and host-derived organosulfonates, such as taurine (2-aminoethansulfonate), as sulfite donors for sulfite respiration, producing hydrogen sulfide (H_2_S), an important intestinal metabolite that may have beneficial as well as detrimental effects on the colonic environment. Its taurine desulfonation pathway involves a glycyl radical enzyme (GRE), isethionate sulfite-lyase (IslAB), which cleaves isethionate (2-hydroxyethane sulfonate) into acetaldehyde and sulfite.

**Results:** We demonstrate that taurine metabolism in *B. wadsworthia* 3.1.6 involves bacterial microcompartments (BMCs). First, we confirmed taurine-inducible production of BMCs by proteomic, transcriptomic and ultra-thin sectioning and electron-microscopical analyses. Then, we isolated BMCs from taurine-grown cells by density-gradient ultracentrifugation and analyzed their composition by proteomics as well as by enzyme assays, which suggested that the GRE IslAB and acetaldehyde dehydrogenase are located inside of the BMCs. Finally, we are discussing the recycling of cofactors in the IslAB-BMCs and a potential shuttling of electrons across the BMC shell by a potential iron-sulfur (FeS) cluster-containing shell protein identified by sequence analysis.

**Conclusions:** We characterized a novel subclass of BMCs and broadened the spectrum of reactions known to take place enclosed in BMCs, which is of biotechnological interest. We also provided more details on the energy metabolism of the opportunistic pathobiont *B. wadsworthia* and on microbial H_2_S production in the human gut.

## Introduction

*Bilophila wadsworthia* is an obligately anaerobic, sulfite-reducing bacterium and part of the normal human gut flora. It has also been associated with diseases such as abscesses, appendicitis, colitis or Parkinson’s disease. *B. wadsworthia* is able to utilize taurine (2-aminoethanesulfonate) as well as isethionate (2-hydroxyethanesulfonate) as sources of sulfite as terminal electron acceptor, in a process termed ‘organosulfonate respiration’ (1), with hydrogen sulfide (H_2_S) as an end product (2). In addition to the H_2_S-production by sulfate-reducing bacteria, organosulfonate respiration may be another important source of H_2_S in the human gut. Bacterial H_2_S production has previously been implicated as an important factor in the development of inflammatory bowel diseases such as colitis (3–5) and colon cancer (6), an effect potentially due to its destructive influence on the mucus barrier of the colon (7, 8) and its genotoxicity (9, 10). On the other side, H_2_S, “the Janus-faced metabolite” (11) can also function as an intracellular antioxidant, a signaling molecule or a mitochondrial energy source, (12) and therefore have beneficial impacts on the host in a dose-dependent manner (13).

Taurine as substrate for *B. wadsworthia* can enter the human digestive system as part of the diet, *e.g*. as a constituent of meat (14, 15) and energy drinks (16), while isethionate can be found in seafood (17, 18). More importantly, taurine is produced by the human body, for example as a conjugate for bile salts such as taurocholate (19–21). It was shown that both a diet rich in saturated fats stimulating bile salt production, including taurocholate, and a diet supplemented directly with taurocholate, can in mice lead to *B. wadsworthia* blooms and subsequent development of colitis (22). The taurine catabolism of *B. wadsworthia* thus provides an intriguing link between dietary conditions, microbial H_2_S production and disease development in animals and humans (21). While *B. wadsworthia* produces H_2_S from organosulfonates, it is unable to catalyze reduction of inorganic sulfate as electron acceptor (2). Its genome lacks the genes for sulfate adenylyltransferase (Sat) and adenosine 5’-phosphosulfate reductase (AprAB), and *B. wadsworthia* is thus unable to activate sulfate and reductively cleave adenosine 5’-phosphosulfate to sulfite and adenosine monophosphate. However, it does contain dissimilatory sulfite reductase complex (Dsr), enabling it to reduce sulfite released from organosulfonates (or free sulfite provided in the culture fluid) further to H_2_S (2).

Only recently, the complete desulfonation pathway for taurine in *B. wadsworthia* was elucidated and shown to include a new class of glycyl radical enzymes (GREs) that cleaves the carbon-sulfur bond in isethionate (1, 23). The complete pathway is illustrated in Figure 1A. Taurine is first deaminated to sulfoacetaldehyde by a taurine-pyruvate aminotransferase (Tpa), which is then reduced to isethionate by a NADH-dependent sulfoacetaldehyde reductase (SarD). Isethionate is the substrate for the GRE isethionate-sulfite lyase (IslA), which is activated by its activating enzyme IslB using S-adenoslymethionine (SAM) as the initial radical donor. This radical-mediated cleavage reaction, which is extremely oxygen sensitive (1), results in the sulfonate group of isethionate being released as sulfite and of the carbon moiety as acetaldehyde. Both sulfite and acetaldehyde can be harmful to cellular processes, as both can damage proteins, DNA and lipids through formation of adducts and are therefore quickly detoxified by most organisms (24, 25).

**Figure 1.**
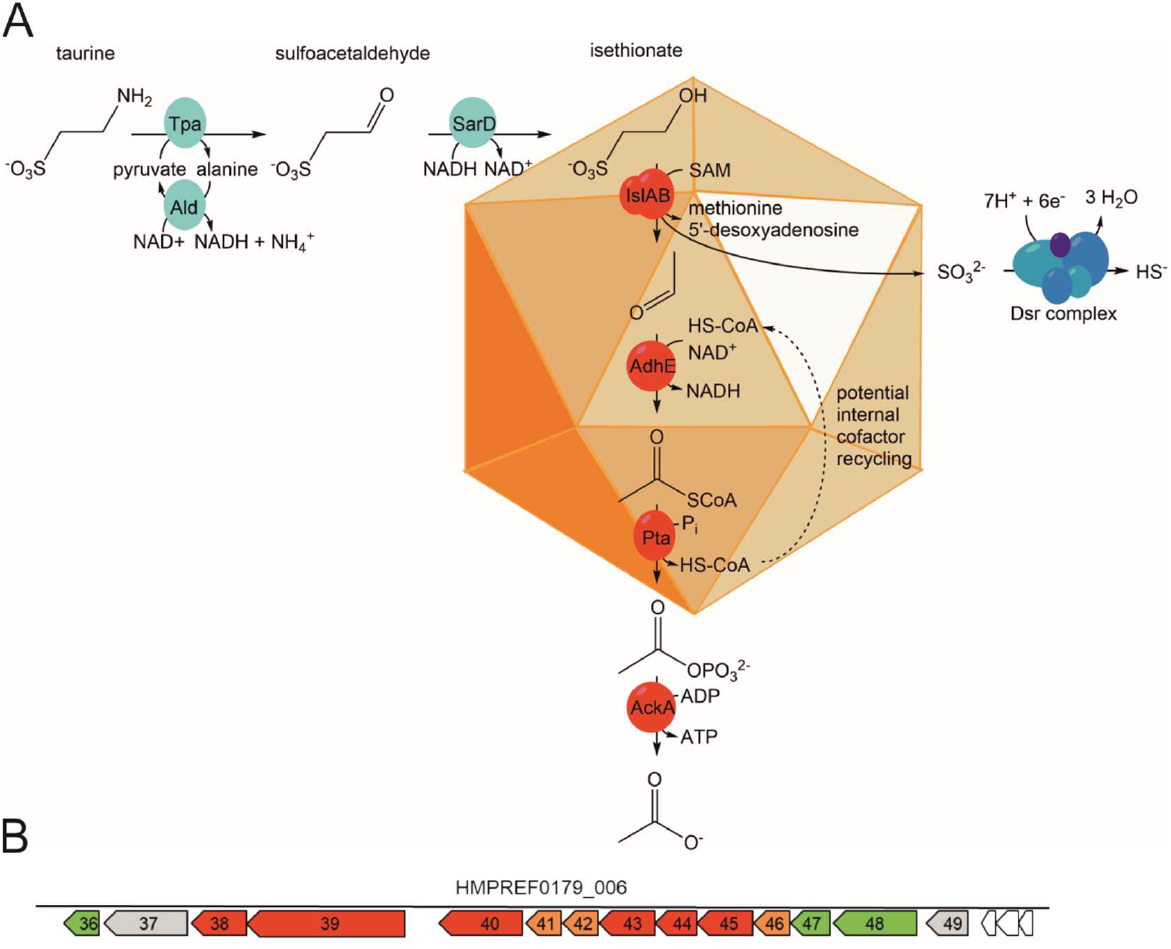
Overview of the taurine desulfonation pathway in *B. wadsworthia* 3.1.6, as revealed previously (1, 26, 27), and of the key enzymes of the pathway enclosed in bacterial microcompartments (BMCs), as inferred from the results of this study. **(A)** Illustration of the taurine degradation pathway *via* isethionate desulfonation by a glycyl radical enzyme and the subsequent conversion of the acetaldehyde released from isethionate to acetate, and of the reduction of the sulfite released to H_2_S by the dissimilatory sulfite reductase complex. The involvement of BMCs in this pathway, as inferred from the results of this study, is also indicated (see main text). Please note that H_2_S, HS^-^ and to a small extent S^2-^ are in equilibrium at physiological pH, but that H_2_S will be used throughout this study to refer to all three species. Enzyme abbreviations used: Tpa, taurine-pyruvate aminotransferase; Ald, alanine dehydrogenase; SarD, sulfoacetaldehyde reductase; IslAB, isethionate-sulfite lyase; AdhE, acetylating acetaldehyde dehydrogenase; Pta, phosphotransacetylase; AckA, acetate kinase; Dsr, dissimilatory sulfite reductase. **(B)** The gene cluster in *B. wadsworthia* 3.1.6 encoding isethionate sulfite-lyase IslAB (Integrated Microbial Genomes (IMG) locus tags HMPREF0179_00638 and _00639) and BMC-associated proteins. Genes marked in red are genes for catabolic enzymes, genes marked in orange encode BMC shell proteins, and genes marked in green may encode proteins that we suspect to be involved in maintaining the redox balance within the BMC (see Discussion). The detailed description for each gene can be found in Table 1.

**Table 1.** Annotations and predicted functions of genes in the IslAB gene cluster. All initial annotations were taken from the original IMG annotation for *Bilophila wadsworthia* 3.1.6. The prefix to yield a complete IMG locus tag is HMPREF0179_.

| Locus tag | Annotation | Predicted gene product function |
| --- | --- | --- |
| 00636 | TIGR04076 family protein | potentially involved in redox reactions or iron cluster metabolism |
| 00637 | PAS domain S-box-containing protein | transcriptional regulator |
| 00638 | pyruvate formate lyase activating enzyme | IslA-activating enzyme, IslB (confirmed, see (1)) |
| 00639 | formate C-acetyltransferase | isethionate sulfite-lyase, IslA (confirmed, see (1)) |
| 00640 | aldehyde dehydrogenase | acetylating acetaldehyde dehydrogenase (confirmed, see (1)) |
| 00641 | ethanolamine utilization protein EutN | BMC shell protein (28, 29) |
| 00642 | ethanolamine utilization protein EutN | BMC shell protein (28, 29) |
| 00643 | phosphotransacetylase | phosphotransacetylase |
| 00644 | ethanolamine utilization cobalamin adenosyltransferase | unclear (see Discussion) |
| 00645 | ethanolamine utilization protein EutQ | putative acetate kinase (30) |
| 00646 | BMC domain-containing shell protein |  |
| 00647 | Carboxysome shell and ethanolamine utilization microcompartment protein CcmL/EutN | protein alignment (BLAST) revealed it to be also similar to PduT, a shell protein from propanediol utilization microcompartments that contain 4Fe-4S cluster (31-33) (see also Discussion) |
| 00648 | Na<sup>+</sup>-translocating ferredoxin:NAD<sup>+</sup> oxidoreductase RNF (RnfC subunit) | based on its four protein family (pfam) domains, we conclude it to be a PduS homolog (see Discussion) |
| 00649 | hypothetical protein | predicted amidohydrolase function (IMG) |

The sulfite released from isethionate is utilized as terminal electron acceptor by the Dsr complex, producing H_2_S (Figure 1A). Notably, this choice of electron acceptor is energetically beneficial compared to sulfate reduction, in that no investment of two ATP equivalents for the activation of sulfate is necessary (1). The acetaldehyde formed (Figure 1A) is oxidized by an acetylating aldehyde dehydrogenase (AdhE) to acetyl-CoA, which is converted to acetyl phosphate by a phosphotransacetylase (Pta) and to, ultimately, acetate after transfer of the phosphate to ADP to form one ATP by an acetate kinase (AckA). The acetate is excreted. As this pathway is not balanced in respect to electrons, an additional electron donor such as lactate, formate or hydrogen (34) is necessary to provide electrons to the Dsr complex for respiration and energy conservation. Indeed, *B. wadsworthia* is able to utilize organosulfonates as terminal electron acceptors but not as sole substrates for fermentation (2, 34).

The genes for the taurine desulfonation pathway are encoded in two clusters that are regulated independently (1): One taurine-inducible cluster (not shown in Figure 1B) encoding for the enzymes for conversion of taurine to isethionate (*i.e*., genes for Tpa, for pyruvate-regenerating alanine dehydrogenase, and for SarD), and a second, taurine- and isethionate-inducible gene cluster (as shown in Figure 1B) comprising the genes for IslA, IslB and AdhE.

Intriguingly, this second, IslAB-AdhE gene cluster comprises also several genes that are predicted to encode for shell proteins of bacterial microcompartments (BMCs) (see Figure 1B), suggesting that BMCs might play a role in the taurine desulfonation pathway in *B. wadsworthia* 3.1.6 (see below). BMCs are small (40-600 nm in diameter; refs. (35-37)) organelle-like compartments constructed entirely from protein that exist within the bacterial cytosol. Their functions are to isolate enzymatically catalyzed reactions, to protect the cell by containing highly reactive and/or volatile intermediates, usually aldehydes, and/or to improve the efficiency of multistep pathways (35). BMCs have first been described in cyanobacteria, in which the CO_2_-fixing ribulose-1,5-bisphosphate carboxylase/oxygenase is contained within the so-called carboxysome (38–41). Later, BMCs were described also in cells catabolizing ethanolamine (28, 42, 43) or propanediol (44) by B_12_-dependent enzymes (45, 46). These catabolic BMCs, also called metabolosomes, are typically defined by their signature enzymes, *e.g*., choline-trimethylamine lyase or B_12_-dependent diol dehydratases. In addition to these aldehyde-forming signature enzymes, metabolosomes usually contain also an aldehyde dehydrogenase that further processes the aldehyde, an alcohol dehydrogenase and a phosphotransacetylase (35). It has been shown that genes encoding BMC shells can be transferred between bacterial species (33, 47, 48) and even between bacteria and plants (49, 50), making them an interesting module for biotechnological engineering. Indeed, BMCs employed as ‘intracellular bioreactors’ can increase bacterial production of ethanol (51) or improve phosphate removal from the medium by increasing the production of polyphosphates (52). Glycyl radical enzymes (GREs) are the signature enzymes of a range of BMCs. Two well-defined examples are choline trimethylamine-lyase (CutC) in, for example, *Desulfovibrio desulfuricans* and *Escherichia coli* (53, 54) and B_12_-independent glycerol dehydratase in *Clostridium butyricum* and *Rhodobacter capsulatus* (55–58). Other classes of potentially GRE-containing BMCs were defined by genomic analyses (59, 60), but as of yet their function has only been studied in one case, for propanediol utilization (58).

In *B. wadsworthia*, the putative genes for BMC shell proteins are co-located in a predicted operon with the genes for IslAB and AdhE (see Figure 1B). Hence, we speculated that the radical cleavage of isethionate by the extremely oxygen-sensitive IslAB, yielding two toxic products, sulfite and acetaldehyde, may be isolated within a BMC in *B. wadsworthia*.

In this study, we demonstrated the production of BMCs in *B. wadsworthia* 3.1.6 and examined their association with the enzymes of the taurine desulfonation pathway. The expression of BMCs in taurine-grown *B. wadsworthia* cells was shown using electron microscope, transcriptomic and proteomic analyses. Further, the BMCs were separated from cell-free extracts using sucrose-gradient centrifugation and examined by electronmicroscopy and proteomic analysis as well as by specific enzyme assays.

## Results

### Transcriptomic and proteomic analyses confirm expression of BMC shell proteins during growth with taurine

The gene cluster (predicted operon) in *B. wadsworthia* 3.1.6 encoding for IslAB and AdhE contains also four predicted genes for BMC shell proteins (Introduction, Figure 1B), and we therefore examined by transcriptomics and proteomics whether these BMC genes/proteins may be co-induced/expressed during degradation of taurine. Taurine-grown cultures of *B. wadsworthia* (n = 3) were analyzed either by proteomics against 3-sulfolactate-grown cultures (n = 3), or by transcriptional analysis (n = 3) against a 2,3-dihydroxypropanesulfonate-grown culture (n = 1) as the reference (see also Discussion). Figure 2A shows a direct comparison of the proteomic and transcriptomic data for relevant proteins/genes as detected each in taurine-grown cells, and Figure 2B shows a comparison of the proteomic data for taurine-grown cells in comparison to 3-sulfolactate-grown cells. The transcriptomic data for the 2,3-dihydroxypropanesulfonate-grown culture is shown in the Supplementary file, Figure S1. Overall, the data confirmed a strong inducible expression of IslAB, AdhE and of the other taurine-degradation genes during growth with taurine (1). The data showed also that the shell protein genes HMPREF0179_00641 and HMPREF0179_00642 (both annotated as EutN), HMPREF0179_00646 (CcmK-like shell protein) and HMPREF0179_00647 (CcmL/PduT), are more strongly induced during growth with taurine as electron acceptor in comparison to 3-sulfolactate, both at the transcriptional (Figure 2A) and at the protein level (Figure 2B). The stronger induction of these genes during taurine degradation was also indicated by transcriptional analysis relative to 2,3-dihydroxypropanesulfonate-grown cells (see Fig. S1). Hence, *B. wadsworthia* 3.1.6 indeed produced BMC shell proteins during taurine degradation and, thus, most likely BMCs. In the next step, we aimed at a visual confirmation of such sub-cellular compartments in *B. wadsworthia* 3.1.6 cells by electron microscopy.

**Figure 2:**
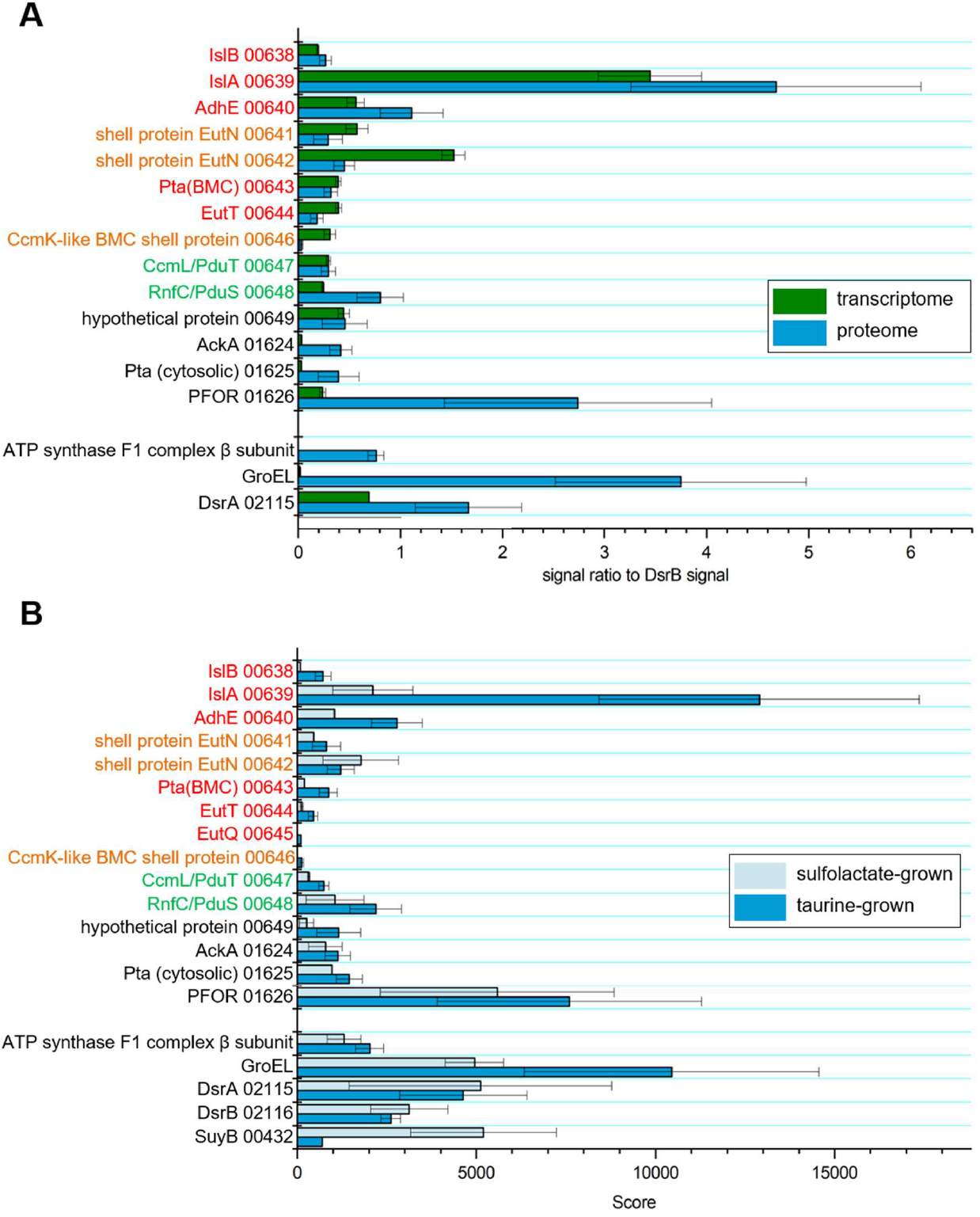
Excerpts of the total transcriptomic and proteomic data for *B. wadsworthia* in respect to the expression of BMC shell proteins during taurine degradation. **(A)**: Proteomic (blue) and transcriptomic (green) identification of produced or induced proteins/genes in *B. wadsworthia* during growth with taurine as electron acceptor. Values are normalized to the DsrB signal to enable comparison between the two methods. Note that the transcription of the genes for the small BMC shell proteins such as (IMG locus tag prefix HMPREF0179_) genes 00641, 00642 (both annotated as EutN) and 00646 (CcmK-like shell protein) appeared to be higher relative to the proteomic scores in comparison to the data for larger genes/proteins (e.g. for IslA, GroEL, PFOR), and that this observation may be attributed to the proteomic analysis, where smaller proteins are detected with lower sensitivity than larger proteins. Error bars: standard deviation of n = 3 **(B)** Excerpt of total proteomic data for *B. wadsworthia* grown with taurine (dark blue) in comparison to 3-sulfolactate (light blue) as electron acceptor. For taurine respiration, a higher expression of IslAB was confirmed, as well as of other BMC-associated proteins from the gene cluster represented in Figure 1B, such as the shell proteins EutN (00641), a CcmK-like shell protein (00646) and CcmL/PduT (00647). Sulfolactate respiration on the other hand led to much higher expression of the corresponding desulfonating enzyme, SuyAB (see text), in comparison to taurine respiration. Data for constitutively expressed proteins is also shown for comparison (e.g. GroEL, ATP synthase subunits). Data represents the mean ± standard error from the analysis of three biological replicates (cultures) for taurine- and of two biological replicates for sulfolactate-grown *B. wadsworthia* 3.1.6.

### Confirmation of BMC production by transmission electron microscopy

Ultrathin sections of epoxy-embedded *B. wadsworthia* cells grown with taurine as the electron acceptor were examined for the existence of BMCs by transmission electron microscopy (TEM) (Figure 3AB). Multiple polyhedral structures of 50-100 nm in diameter were visible, essentially, in all cells examined (see Figure 3A), resembling BMCs when compared against the BMC structures in *Desulfovibrio alaskensis* cells grown with choline (Figure S2), hence, under conditions that are known to lead to BMC formation in this organism (61). These structures were not present in *B. wadsworthia* cells grown with 3-sulfolactate (Figure 3B). 3-sulfolactate is desulfonated by a different inducible enzyme, 3-sulfolactate sulfite-lyase (SuyAB) (62, 63) (see Figure 2B). SuyAB is not oxygen sensitive and produces sulfite and non-toxic pyruvate, in comparison to the acetaldehyde produced by IslAB (see the Discussion) and is therefore not expected to lead to the formation of BMCs. Further, the gene cluster for SuyAB does not contain BMC shell protein genes (Figure S3). Correspondingly, the shell proteins encoded in the IslAB gene cluster were detected only at low abundance in cells grown with 3-sulfolactate (Figure 2B).

**Figure 3:**
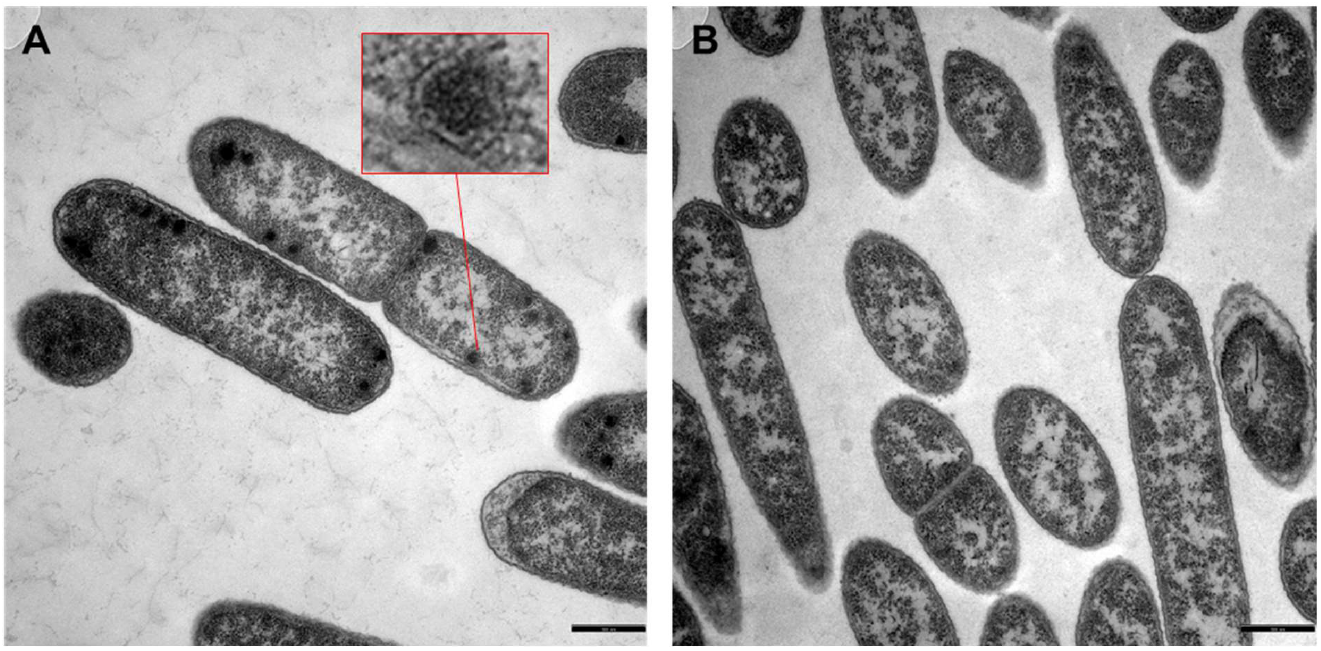
TEM images of ultrathin sections of *B. wadsworthia* cells grown with taurine (A) or 3-sulfolactate as the electron acceptor (B). Polyhedral structures characteristic of BMCs were observed specifically for taurine-grown cells. Scale bars: 500 nm. The inset in (A) is a 6x magnification. The structures in *B. wadsworthia* appeared very similar to these in the BMC-producing *Desulfovibrio alaskensis* during growth with choline (see Figure S2), as previously described (64).

### Separation of BMCs by sucrose gradient centrifugation

We aimed at purifying the BMCs using a sucrose gradient centrifugation, which separates cell components according to their sedimentation speed (see Material and methods). As shown in Figure 4, a total of ten clearly distinguishable bands were visible after centrifugation and were collected separately into ten fractions. SDS-PAGE analysis was employed for preliminary attribution of these fractions (Figure 4C): The distinctive band of the glycyl radical enzyme IslA (93.9 kDa molecular weight, confirmed by proteomic analysis and marked with the red box in Figure 4C) was found in fractions 3 to 5, and was most prominent in fraction 4, suggesting that this was the main fraction in which also BMCs may be contained. Indeed, when fraction 4 was analyzed by TEM, it was found to contain microcompartment-like structures (Figure 4B), suggesting that also intact BMCs were purified in fraction 4. Fraction 2 contained a diverse mixture of proteins, but no distinct band at 94 kDa, and we presumed this to be the fraction of predominantly soluble cytosolic proteins.

**Figure 4:**
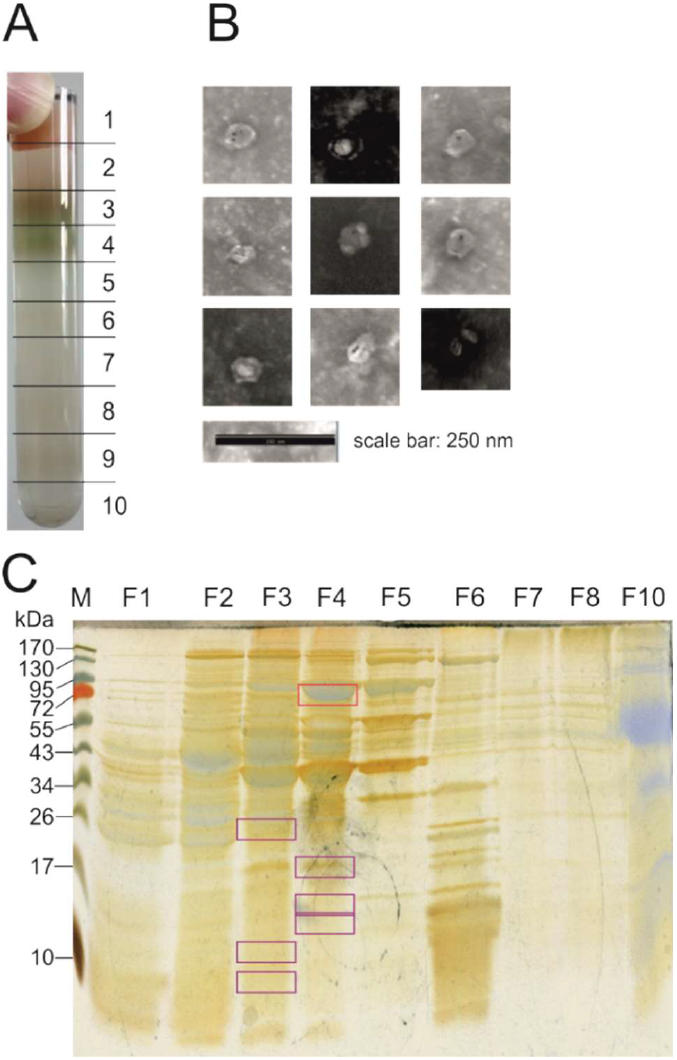
Sucrose gradient purification of BMCs from cell-free extracts of *B. wadsworthia*. **(A)** Photograph of a sucrose gradient after centrifugation. The numbers mark the fractions that were collected. Fraction 2 contained most of the soluble cytosolic proteins, fraction 3 broken BMCs, while fraction 4 contained intact BMCs (see Figure 4C). Fraction 10 contained also the centrifugation pellet; it was resuspended in fraction-10 solution (0.5 ml) and included in the SDS-PAGE analysis. **(B)** Negative-stained TEM images of BMCs as visible in fraction 4. **(C)** Double-stained (Coomassie and silver) SDS-PAGE gel of the gradient fractions. The upper red box marks the molecular weight of the signature enzyme of the BMC, isethionate sulfite-lyase (IslA) at approximately 94 kDa. The strong band at 40 kDa correspond to SarD (40.8 kDa) and Ald (39.8 kDa). The purple boxes mark bands that were identified by proteomic analysis as BMC shell proteins for fractions 4 and fraction 3 (see main text), which were visible only through silver staining. Fraction 9 was not included in the analysis as it contained only an insignificant amount of protein. M, protein molecular weight markers.

The SDS-PAGE bands from fractions 3 and 4 in the size range of predicted BMC shell proteins were cut out from the gel and analyzed by proteomics (Table S2). These bands were identified as EutN (HMPREF0179_00642 and HMPREF0179_00647), a known shell protein from ethanolamine utilization BMCs, among many other proteins, especially ribosomal proteins (see Table S2). We concluded that the fractions containing IslA also contained identified BMC shell proteins, and that ribosomes, in size and composition comparable to small BMCs, may have been co-purified with the BMCs under the conditions were used.

### Proteomic analysis of the sucrose gradient fractions

The BMC-containing fraction 4 was analyzed also by semi-quantitative total proteomics (*i.e*., without SDS-PAGE separation) relative to fractions 2, 3 and 5. Figure 5 shows the relative abundance of selected proteins within each dataset and throughout the gradient fractions (yellow, fraction 2; orange, fraction 3; red, fraction 4 [BMC fraction]; pink, fraction 5).. The cytosolic chaperone DnaK served as a control. It was detected in higher relative abundance (*i.e*., with higher score) in fractions 2 and 3 than in fractions 4 and 5, as expected for a cytosolic protein. For the components of the BMC signature enzyme IslAB a contrasting distribution was found. They were more prominently detected in fractions 4 and 5, in which based on the results described above intact BMCs were observed by TEM (Fig. 4B) and in which BMC shell-proteins and IslA were identified after SDS-PAGE separation (Fig. 4C). This further confirmed that fractions 4 and 5 were the BMC-containing fractions. Interestingly, a similar abundance pattern was observed for the DsrAB components of sulfite reductase, for pyruvate:ferredoxin oxidoreductase (PFOR) and to a certain extent also for aldehyde dehydrogenase (AdhE), phosphotransacetylase (Pta), the RnfC subunit oxidoreductase and alanine dehydrogenase (Ald). Shell proteins, acetate kinase (AckA), and the enzymes converting taurine to isethionate (Tpa, SarD), appeared to be more abundant in the cytosolic fraction 2 and in the intermediate fraction 3. The shell proteins detected in the soluble protein fractions resulted most likely from fractured BMCs or monomeric shell proteins (10-20 kDa).

**Figure 5:**
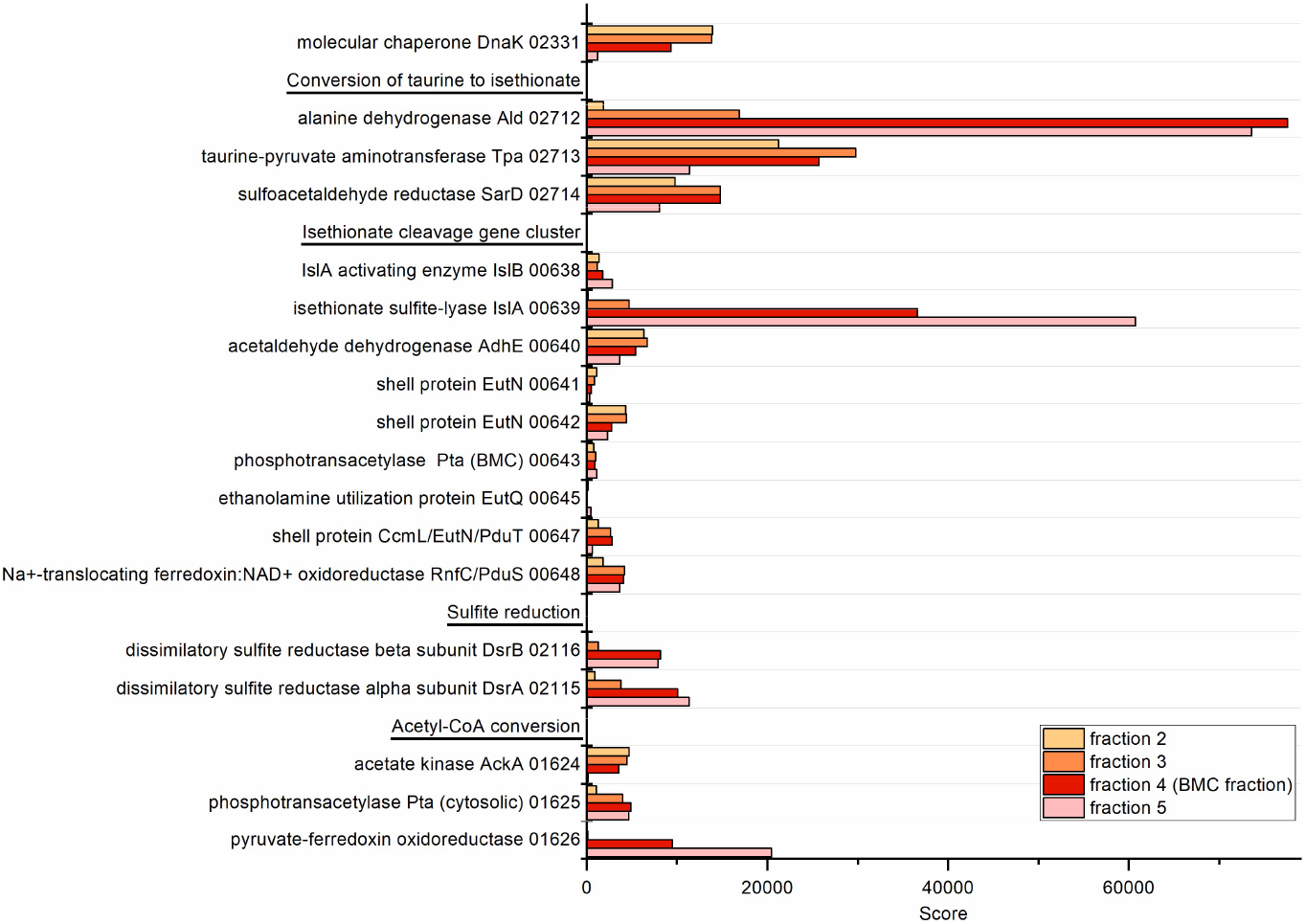
Excerpt of the total proteomic data obtained for the sucrose gradient fractions. Legend: yellow, fraction 2; orange, fraction 3; red, fraction 4 (BMC fraction); pink, fraction 5. Protein numbers refer to IMG locus tags without prefix (HMPREF0179_) as identifiers.

### Specific enzyme activities detected in sucrose gradient fractions

We also measured specific activities of the taurine pathway enzymes associated to the different gradient fractions (Figure 6), aiming to relate these activities to the protein abundances as observed by proteomics.

**Figure 6:**
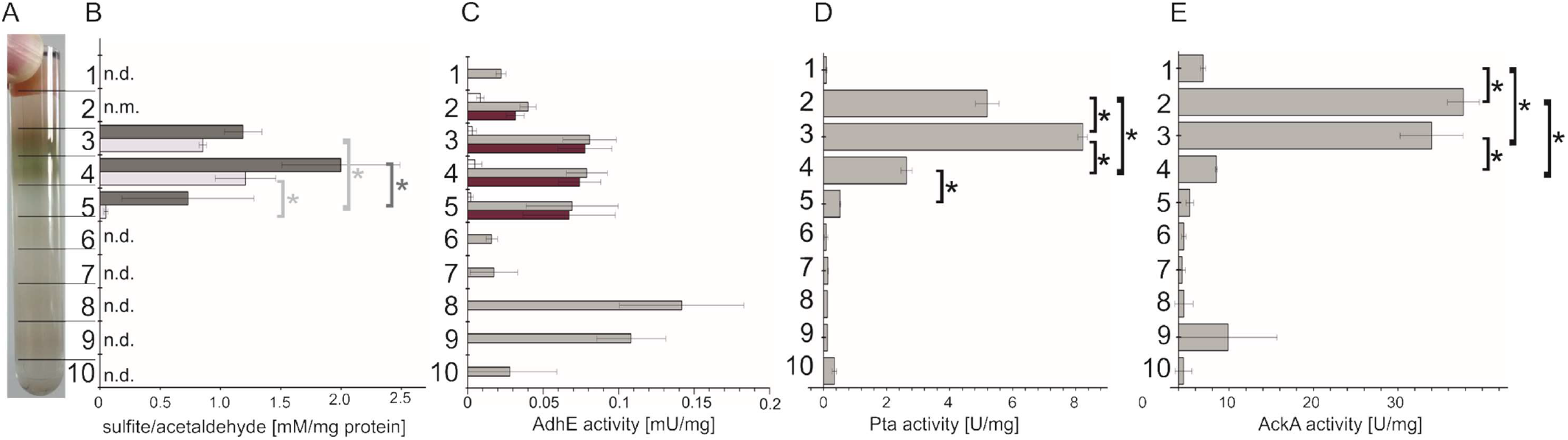
Specific enzyme activities detected in the different sucrose gradient fractions (B-E) in comparison to an image of a representative gradient (A). Protein concentrations in the fractions 7-9 were typically very low and, thus, apparently high specific enzyme activities detected were considered unreliable. Specific protein activities are given as Units per mg of total protein. The figure shows means ± standard deviation of three technical replicates, representing one of two biological replicates. Asterisks denote statistical significance at a power of 0.05 **(B)** IslAB activity represented as sulfite (light gray bars) and acetaldehyde (dark gray bars) production in anoxic incubations with isethionate and S-adenosylmethionine. n.d.: not determined; n.m.: not measurable. **(C)** Specific acetaldehyde dehydrogenase activity determined photometrically as rate of NADH formation from NAD^+^. Grey bars, total acetaldehyde dehydrogenase activity (measured in presence of free coenzyme A); red bars, acetylating acetaldehyde dehydrogenase activity, calculated by subtracting non-acetylating acetaldehyde dehydrogenase activity (measured in absence of free coenzyme A). **(D)** Specific phosphotransacetylase activity measured as acetyl-CoA formation from acetyl phosphate in the presence of free coenzyme A. **(E)** Specific acetate kinase activity measured as acetyl phosphate consumption in the presence of ADP.

Isethionate-cleaving activity of IslAB was measurable in the BMC fraction 4 and to a lower extent in the neighboring fractions 3 and 5, confirming the distribution displayed by the proteomic data (Figure 6B). Both the formation of acetaldehyde and sulfite was detectable in the BMC fractions after incubation with isethionate and S-adenosylmethionine in assays under strictly oxygen-free, reducing conditions (see Material and Methods). For comparison, there was no acetaldehyde or sulfite formation measurable in the soluble protein fraction 2.

The detected acetylating aldehyde dehydrogenase activity was evenly distributed across the fractions 3 to 5, again confirming the distribution pattern found in the proteomic data (Figure 6C). Almost all of the aldehyde-oxidizing activity was found to be coenzyme A-dependent, while non-acetylating acetaldehyde oxidation was negligible.

Phosphotransacetylase activity was detected at the highest level in the intermediate fraction 3 and slightly lower in fractions 2 and 4 (Figure 6D). As discussed below, *B. wadsworthia* encodes two phosphotransacetylases in its genome, which cannot be distinguished by the activity measurement; they both might have been present in different distribution along the gradients.

The highest acetate kinase activity was detected in the soluble fractions 2 and 3, confirming its distribution as suggested by the proteomic data (Figure 6E). The activity was markedly lower in the BMC-containing fractions 4 and 5. That this activity pattern of acetate kinase was different to that of phosphotransacetylase, which seemed to be more closely associated to the BMCs, is consistent with the distinct distribution pattern as displayed by the proteomics data.

The activity of taurine-pyruvate aminotransferase (Tpa) was not measurable as the enzyme did not remain active throughout the centrifugation. Further, it was impossible to determine sulfoacetaldehyde reductase (SarD) activity in a photometrical assay against a background NADH-oxidizing activity, but previous work with recombinant SarD (published in (1)) suggests that it would retain its full activity throughout the centrifugation process.

## Discussion

In this study, the existence of BMCs in *B. wadsworthia* during growth with taurine as electron acceptor, was demonstrated by total-proteomic and transcriptomic analyses as well as by electron microscopy. Further, we succeeded in enriching the BMCs by gradient centrifugation, as confirmed by electron microscopy and proteomic analysis, and we were able to confirm IslAB enzyme activity and its proteins being associated to the BMC-containing fraction. Based on the clear distribution patterns of IslAB found in both the omics data and activity measurements, we conclude that IslAB is indeed the signature enzyme of these BMCs. *B. wadsworthia* likely employs these BMCs in order to isolate the desulfonating glycyl radical enzyme IslAB, which cleaves isethionate to harmful acetaldehyde and sulfite, from the cytosol. Furthermore, enzymes that process these toxic reaction products of IslAB were found to be associated to the BMCs, though it remains unclear whether they are contained within, or are merely associated to the outside of the BMC, such as the aldehyde-detoxifying dehydrogenase AdhE (discussed further below) and the sulfite-reducing and, thus, sulfite-detoxifying Dsr complex, of which the DsrA and B components also seemed to be associated to the BMC-containing fraction (Fig. 5). Whether the Dsr complex may be associated to, most likely, the outside of the BMCs for an efficient detoxification of sulfite remains to be addressed in future research. The TEM images of the ultra-thin sections of taurine-grown *B. wadsworthia* (Fig. 2A) suggest for many cells a ‘crowding’ of cellular structure particularly around the BMCs and, hence, it is tempting to speculate whether the BMCs may indeed be covered by densely packed peripheral enzymes, such as in this case, maybe the Dsr complex.

Recently, another desulfonating GRE was demonstrated in *B. wadsworthia*, 2,3-dihydroxypropanesulfonate (DHPS) sulfite-lyase (HpsGH), for growth with DHPS as sulfite-donor for respiration (11, 65) (see Fig. S1). This GRE is not encoded in a gene cluster associated with BMC shell protein genes and, thus, most likely is a free, cytosolic GRE in *B. wadsworthia*. The reaction product hydroxyacetone from desulfonation of DHPS might be less toxic compared to the acetaldehyde formed during the IslAB reaction. However, *Desulfovibrio desulfuricans* DSM642 and *Desulfovibrio alaskensis* G20 also employ IslAB-type GREs for isethionate desulfonation (1), and these enzyme systems are expressed from gene clusters that also do not co-encode BMC shell proteins. Hence, there seems to be no absolute necessity to encapsulate the IslAB reaction in BMCs for protection against toxicity of sulfite and acetaldehyde. *Desulfovibrio*, as ‘classical’ sulfate reducing bacteria being able to utilize sulfate, may utilize organosulfonates as alternative sulfite donor for respiration, while *B. wadsworthia* has specialized solely on organosulfonate respiration. The encapsulation of the IslAB-GRE reaction in BMCs might in *B. wadsworthia* thus reflect a structural specialization for competitive advantage.

The activity of acetylating acetaldehyde dehydrogenase could clearly be associated to the BMCs of *B. wadsworthia*. As known from other types of BMC (35) that harbor aldehyde dehydrogenases, acetylating acetaldehyde dehydrogenase contained within the BMC would be keeping concentrations of free acetaldehyde low. The acetyl-CoA resulting from the oxidation of acetaldehyde would be trans-esterified to acetylphosphate by Pta, a protein that, based on our proteomic data and activity measurements, seemed to be associated with the BMC, and then converted to acetate and ATP by cytosolic acetate kinase. As our enzyme assay data shows high Pta activity also in the BMC fraction, transformation of acetaldehyde *via* acetyl-CoA to acetylphosphate in the BMC is probable. The acetylphosphate would likely be shuttled out of the BMC and dephosphorylated to acetate by AckA (coupled to ATP formation), which is, based on the enzymes assays, most probably a cytosolic enzyme.

In this case, for acetylating acetaldehyde dehydrogenase and Pta being located within the BMC, coenzyme A would be regenerated within the BMC. As coenzyme A is a large cofactor compared to, for example, acetylphosphate, regeneration within the BMC would save substantial transport across the shell. Such internal cofactor recycling has been shown for ethanolamine- (66) and 1,2-propanediol-utilization (67) microcompartments. For ethanolamine utilization of *Salmonella enterica*, the gene for the phosphotransacetylase EutD in the BMC gene cluster has been shown by Huseby and Roth (66) to be essential even though it is functionally redundant since a second, housekeeping phosphotransacetylase gene exists elsewhere in the genome. The authors assumed that the BMC-contained Pta is essential due to its ability to recycle free coenzyme A within the BMC. A similar situation may be the case in *B. wadsworthia*, which was found to express two phosphotransacetylases as well: One whose respective gene is located within the BMC gene cluster (HMPREF0179_00643) and one that is encoded next to cytosolic acetate kinase (HMPREF0179_01625). Both of these phosphotransacetylases were found associated with the BMC by proteomic analysis; however, the assay we used cannot distinguish between the activities of these two enzyme homologs. It is possible that one is indeed located within the BMC and one in the cytosol. The BMC gene cluster even contains a gene annotated as EutQ which has been described as having an acetate kinase activity (30) and could catalyze the last reaction step from acetyl phosphate to acetate, but it was only detectable in low amounts in proteomic analysis; in addition, the acetate kinase activity was mainly detected in the soluble protein fraction.

The potential internal recycling of NAD^+^ from NADH formed by acetaldehyde oxidation to acetyl-CoA is much more unclear, though an intriguing candidate may be a protein annotated as Na^+^-translocating ferredoxin:NAD^+^ oxidoreductase (RnfC, 00648). In our proteomic analysis, this protein was found associated with the BMCs, for which it could serve as part of a redox shuttle in order to remove excess electrons from the BMC across the shell, thereby internally regenerating NAD^+^ from NADH. Based on the protein-family (pfam) domains, of which the motif of the RnfC subunit (pfam13375) is only one of four (68), it is more similar to the cobalamin reductase PduS from *Salmonella enterica* or *Citrobacter freundii* than to the actual Rnf complex subunit. PduS in propanediol utilization microcompartments reduces the cobalamin cofactor of B_12_ from Co^3+^ to Co^2+^ (69, 70) and also associates with PduT, a shell protein containing a 4Fe-4S cluster (70). A PduT homolog may be encoded in the *B. wadsworthia* gene cluster annotated as CcmL/EutN (00647): Basic Local Alignment Search (BLAST) search revealed the most closely related protein (69.7% identity) to be a protein of the PduT family in *Telmatospirillum siberiense*. Its identity to the experimentally verified PduT sequences of *C. freundii* and *Salmonella typhimurium* strain LT2 is 43.9 and 41.2% respectively. In the GRE-containing BMCs of *B. wadsworthia*, which do not involve B_12_ as a cofactor, the function of PduS and PduT is yet unclear. However, it is tempting to speculate that they may also serve to shuttle electrons between the outside and the inside of the BMC using the FeS cluster of PduT, as it was suggested for propanediol (68, 70) and ethanolamine (71) utilization microcompartments. Since the *B. wadsworthia* gene cluster contains no gene for an alcohol dehydrogenase in order to recycle NAD^+^ within the BMCs (66, 67), it remains unclear how the electrons released by the acetaldehyde dehydrogenase are being shuttled further. Hence, a potential transfer of these electrons to the cytosol *via* the iron-sulfur clusters of PduS in the BMC lumen and PduT in the BMC shell is a viable hypothesis. It has also recently been shown that in *Listeria monocytogenes*, ethanolamine is anaerobically catabolized in BMCs but at the same time dependent on extracellular electron transport (71), necessitating a transfer of electrons out of the BMC. In this study, the authors also suspect this transport to take place *via* a PduT-like shell protein.

Both PduS and PduT are known to use flavins as well as iron-sulfur clusters as electron carriers. The electron potential of the reduced flavin cofactor of oxically purified recombinant PduS is at −262 mV, in anoxically produced PduS, binding both the flavin cofactor and an [4Fe-4S] center it is at −150 mV (70). This means that the electrons could well be used for sulfite reduction (−116 mV). However, the potential of electrons on the [4Fe-4S] cluster of PduT, which is the proposed redox link in the BMC shell, is +99 mV (33) and not sufficient for sulfite reduction.

Inside the BMC, these flavins could additionally be used to shuttle electrons to IslB, which like other GRE activating enzymes might require electrons transferred by flavins to install the glycyl radical on IslA (72–74). However, once installed, the radical is regenerated after each reaction of IslA without further electron input, meaning that IslB cannot take up the stoichiometric amounts of electrons produced by AdhE. This makes a second electron sink necessary, such as a transport out of the BMC, for example, to pyruvate:ferredoxin oxidoreductase or DsrAB. The detailed function of apparently B_12_-regenerating enzymes in BMCs with B_12_-*independent* signature enzymes in combination with the unclear redox balance of the BMC, is an exciting field for further study and could provide valuable insight into the very specialized metabolism of the human gut bacterium *B. wadsworthia*. BMCs in propanediol utilization of pathogenic *Salmonella enterica* serovar Typhimurium have been shown to be a fitness advantage for the pathogen (75, 76), meaning that also in the opportunistic pathogen *B. wadsworthia*, a medical relevance is possible.

## Conclusions

This work characterized a novel subclass of BMCs involving a carbon-sulfur bond-cleaving GRE as the signature enzyme. Aside from shedding light on the energy metabolism of the common human-gut symbiont and pathogen *B. wadsworthia*, these results improve our knowledge of the biochemical reactions that can take place within BMCs. As BMCs are of interest for biotechnological applications such as production of chemicals or removal of contaminants from water (51, 52), this knowledge can open doors for an even broader application of their potential.

## Methods

### Growth and harvest of *B. wadsworthia*

*B. wadsworthia* 3.1.6 was grown in anoxic, carbonate-buffered, Ti(III)-reduced minimal medium as described before (1). The medium additionally contained 200 μg/l naphthochinone; 20 mM lactate and 20 mM taurine were supplied as the electron donor and acceptor, respectively. The cells were harvested by centrifugation after 20 hours of growth and the cell pellets stored at −20 °C. For BMC purification, the cells were opened by three passages through a cooled French pressure cell and then centrifuged for 10 minutes at 16 000 x g to remove cell debris. *Desulfovibrio alaskensis* G20 was grown as a positive control for BMC formation in the same medium without naphtochinone and supplied with 40 mM choline as sole substrate (53).

### Transcriptomics

For RNA isolation, 50 ml of overnight culture were centrifuged and the pellet was washed with 1 ml RNA*later*™ (Invitrogen/Thermo Fisher Scientific, Waltham, MA, USA), then resuspended in 100 μl RNA*later*™ and stored at −80 °C. RNA was isolated using a *Quick*-RNA Miniprep kit (Zymo Research, Irvine, CA, USA) and DNA removed using a TURBO DNA-*free*™ kit (Invitrogen/Thermo Fisher Scientific, Waltham, MA, USA). PCR using 16S forward and reverse primers was performed to confirm the absence of DNA contamination. Samples were prepared from triplicates of *B. wadsworthia* cells grown with lactate as carbon source and taurine or dihydroxypropanesulfonate as electron acceptor.

Libraries were constructed at Eurofins Genomics (Ebersberg/Konstanz, Germany) after ribosomal RNA depletion and cDNA synthesis using random hexamer primers and sequenced single-end 50bp on an Illumina HiSeq4000 platform.

For data analysis, the reference genome and annotation file for *Bilophila wadsworthia* 3_1_6 were obtained from NCBI (reference sequence: NZ_ADCP00000000.2). Raw sequence reads were quality trimmed and adapters were removed using the program Trimmomatic 0.39 (77) using the following parameters: minimum read length 30 bp, minimum average phred quality must be at least 15 in a sliding window of 4, bases at the start and end of sequences are removed if the phred quality is lower than 3. Only a small amount of reads (< 0.5 %) were dropped this way (Table S1). The trimmed reads were aligned to the reference genome using the program Bowtie2 (version 2.3.5.1; (78)) with default settings and the program Samtools (version 1.9 (79)) was used to convert mapping files from “sam” to “bam” format and sort them by genomic coordinates. Stringtie (version 1.3.5 (80)) was used to assign read counts to each gene present in the provided annotation, and in combination with the PERL script *prepDE.pl* (http://ccb.jhu.edu/software/stringtie/) the raw count table (Table S1) was obtained as well as the transcripts per kilobase million (TPM) for each individual gene (Figure S2). For comparison to proteimic data, these values were normalized to the DsrB gene transcript count.

### Sucrose gradient centrifugation

The sucrose gradient was prepared with solutions containing 10, 20, 30, 40, and 50 % (w/v) sucrose in 50 mM Tris-HCl at pH 7.4. 2.5 ml of each solution were used to form the gradient in a 14 ml Ultra-Clear™ centrifuge tube (Beckman Coulter, Brea, CA, USA), with the highest sucrose concentration at the bottom of the tube. The cell-free extract was layered on top and the gradients were centrifuged at 85365 x *g* (average relative centrifugal force in a Beckman Coulter SW 40 Ti swing bucket rotor) for 16 hours at 4 °C (81). The resulting gradient fractions were collected in the same pattern for every individual gradient.

For proteomic analysis, the buffer was exchanged to 50 mM Tris-HCl (pH 7.9) without sucrose using 10 kDa mass cutoff centrifugal filters (Sartorius Vivaspin 500, Sartorius AG, Göttingen, Germany), the samples were then brought to comparable protein concentrations and submitted to the Proteomics Facility of the University of Konstanz for protein mass fingerprinting. For electron a 5 mM Tris -HCl (pH 7.9) without sucrose was used.

### SDS-PAGE and enzyme identification

The protein content of the individual fractions was determined by Bradford assay in a total volume of 1 ml (82). For SDS-PAGE, gradient fractions corresponding to 50 μg of protein were boiled with a loading dye (Roti^®^-Load 1, Carl Roth, Karlsruhe, Germany) and loaded onto a gel (12 % acrylamide in the resolving gel, 4 % in the stacking gel). The gels were run at 100 V for 1 hour and the proteins were visualized by Coomassie and/or silver staining (Table S3).

Individual bands were cut out of the gel and submitted for protein mass fingerprinting at the Proteomics Facility of the University of Konstanz.

### Transmission electron microscopy

#### Negative staining of isolated BMCs

Glow discharge carbon-coated nickel grids were cleaned by oxygen plasma for 45 seconds in a Harrick PDC-32G2 PlasmaCleaner (Harrick Plasma, Ithaca NY, USA). 15 μl sample were applied to the grid by floating the grid on top of the sample for 2 minutes. The grid was then rinsed four times with double-distilled water and contrasted with 1% phosphotungstenic acid neutralized with sodium hydroxide (5 second rinse followed by 45 second incubation in the contrasting solution). The grids were carefully blotted from the edge and dried thoroughly before imaging.

#### Embedding and ultrathin sectioning of cells

Culture samples of *B. wadsworthia* grown with taurine and lactate, of *B. wadsworthia* grown with sulfolactate and lactate and of *D. alaskensis* grown on choline were submitted to the Electron Microscopy Centre at the University of Konstanz.

The cells were enclosed in agarose and fixed with 2.5% glutardialdehyde in 0.05M HEPES buffer at pH 7 for 2.5 hours. After initial dehydration in a graded ethanol series, postfixation was performed with 2 % OsO_4_ for one hour at 0°C, and samples were stained en-bloc with a saturated solution of uranyl acetate in 70% ethanol. Following further dehydration in a graded acetone series the agarose blocs were embedded in Spurr’s resin (Spurr’s Low Viscosity embedding kit, Sigma Aldrich, St Louis, USA) using acetone as intermedium, and polymerized at 65 °C for 48 hours. A detailed description of the process can be found in Table S4.

Ultramicrotomy was performed on a Leica UC7 ultramicrotome (Leica Microsystems, Wetzlar, Germany) using a diamond knife (Diatome 45°Ultra, diatome, Nidau, Switzerland). Ultrathin sections (50nm) were mounted on formvar-coated copper mesh grids, incubated on a drop of saturated aequous uranylacetate solution (~ 7.8%) for 20 minutes in the dark, then rinsed with double distilled water for 10 seconds and dried. They were then incubated for 90 seconds on a drop of 0.4% lead citrate and 0.4% sodium hydroxide solution under carbon dioxide-free conditions, rinsed for 10 seconds with double distilled water and dried overnight under air according to (83).

#### Transmission electron microscopy imaging

Samples were analyzed using a Zeiss EM 912 Omega (Zeiss, Oberkochen, Germany) transmission electron microscope equipped with a thermionic tungsten cathode, and a TRS slow scan CCD-camera for TEM (Tröndle Restlichtverstärker Systeme, Moorenweis, Germany). The images were taken at an acceleration voltage of 80 kV and processed using ImageSP ver.1.2.3.36 (SYS-PROG, Minsk, Belarus & Tröndle Restlichtverstärker Systeme, Moorenweis, Germany).

### Enzyme assays

#### Taurine-pyruvate aminotransferase (Tpa)

Taurine-pyruvate aminotransferase activity was measured aerobically as taurine consumption and alanine formation in a solution of 5 mM taurine and 5 mM pyruvate in 50 mM Tris -HCl at pH 7.9. The cofactor pyridoxal-5-phosphate was additionally supplied in a concentration of 0.1 mM in all buffers used in the assay or during purification. Typically, 0.2 ml of cell-free extract or gradient fraction were used for an assay on a 1 ml scale. Taurine and alanine concentrations were measured by HPLC as described in (1). Each fraction was assayed in triplicates.

#### Sulfoacetaldehyde reductase (SarD)

SarD activity was measured under oxic conditions in 50 mM Tris -HCl buffer at pH 7.9 with 0.4 mM sulfoacetaldehyde and 0.2 mM NADH provided as substrates. The decrease of NADH concentration was measured at 340 nm in a spectrophotometer (JASCO, Tokyo, Japan). 100 μl of gradient fraction were used in one assay and the background activity of the protein solution with NADH only was measured separately before adding sulfoacetaldehyde. Each fraction was assayed in triplicates.

#### Isethionate sulfite lyase (IslAB)

For measuring isethionate sulfite lyase activity, the gradient was prepared from anoxic sucrose solutions under a 95 % N_2_/5 %H_2_ atmosphere and centrifuged in sealed tubes as described above. The fractions were separated in a 95 % N_2_/5 %H_2_ atmosphere. The enzyme assays were carried out in anoxic 50 mM Tris-HCl buffer at pH 7.9. The reaction mixture contained 20 mM isethionate, 1 mM SAM hydrochloride, 2 mg/l resazurin as a redox indicator and 1 mM Ti(III)-NTA as a reducing agent. The reactions were carried out in anoxic stoppered 1 ml glass cuvettes with a 95 % N_2_/5 %H_2_ gas phase. The reaction was started by addition of 200 μl/ml gradient fraction solution. Samples were taken at one hour intervals using analytical syringes. Each fraction was assayed in triplicates.

For acetaldehyde quantification, 50 μl of the reaction were added to a 2.5 mM solution of 2,4-dinitrophenylhydrazine (DNPH) in acetonitrile with 0.1 % phosphoric acid. The derivatization reaction was incubated for at least 30 minutes at room temperature (modified after (84)).

For sulfite quantification, 50 μl of the reaction were added to 0.25 M borate buffer with 0.25 M KCl and 20 mM EDTA, adjusted to pH 10 with sodium carbonate. 50 μl of a 10 mM solution of *N*-(9-acridinyl)-maleimide (NAM) in acetone were added and the reaction was incubated for 30 minutes at 50 °C (85).

All derivatized samples were frozen once to precipitate proteins and then centrifuged for 30 seconds. Derivatized sulfite respectively acetaldehyde were analyzed by HPLC on a phenomenex Luna Omega C18 column (5 μm PS, 100 Å, 150 x 3 mm, phenomenex/Danaher; Washington D.C., USA) with acetonitrile and 0.1 % formic acid in water as mobile phases. The Shimadzu Prominence HPLC (Shimadzu, Kyoto, Japan) was equipped with a SPD-M20A photodiode array detector and the detection wavelengths were 254 nm for NAM-sulfite adducts and 360 nm for DNPH derivatives.

The HPLC gradient programmes were as follows: For DNPH derivates: 25 % acetonitrile for 4.5 minutes, followed by a rise to 70 % acetonitrile over 15 minutes and 10 minutes reequilibration to 25 % acetonitrile. For NAM-sulfite: 3 minutes at 10 % acetonitrile, gradient to 80 % acetonitrile during 10 minutes, reequilibration to 10 % acetonitrile for 7 minutes.

#### Acetylating acetaldehyde dehydrogenase (AdhE)

To determine acetaldehyde dehydrogenase activity, the reduction of NAD^+^ to NADH was measured photometrically (modified after (86)) under oxic conditions. One assay typically contained 0.25 mM NAD^+^, 5-25 μl gradient fraction, 2 mM acetaldehyde and 0.33 mM coenzyme A in a total volume of 1 ml anoxic 50 mM potassium phosphate buffer with 3 mM dithiothreitol (DTT) at a pH of 7.5. The formation of NADH was monitored for one minute at 340 nm. To be able to calculate the activity of acetylating acetaldehyde dehydrogenase specifically, each fraction was measured in one assay with coenzyme A, yielding total acetaldehyde dehydrogenase activity, and one without coenzyme A to measure non-acetylating acetaldehyde dehydrogenase activity. The activity of acetylating acetaldehyde dehydrogenase was calculated as the difference between these two measurements. Each fraction was assayed in triplicates.

#### Phosphotransacetylase (Pta)

Phosphotransacetylase activity was measured as the backwards reaction from acetyl phosphate to acetyl-CoA, with the acetyl-CoA formation being measured photometrically ((87) modified after (88)). 3.33 mM acetyl phosphate and 0.33 mM coenzyme A were incubated i 50 mM Tris -HCl buffer at a pH of 7.5. 10 to100μl of gradient fraction per ml were added and the formation of acetyl-CoA was monitored for one minute at 233 nm. Each fraction was measured in triplicates.

#### Acetate kinase (AckA)

The activity of acetate kinase was measured as the consumption of acetyl phosphate, which was quantified using a colorimetric assay based on the formation of acyl-hydroxamate chelation with Fe^3+^ ((87) modified after (89)). The enzymatic reaction was set up in a volume of 1 ml containing 5 mM ADP, 5 mM MgCl_2_, 3.33 mM acetyl phosphate and 5 to 25 μl gradient fraction in 50 mM Tris -HCl buffer at a pH of 7.4. The reactions were incubated shaking at room temperature (23 °C) and samples of 300 μl for the determination of acetyl phosphate concentration were taken at time intervals of 2 or 5 minutes depending on the reaction speed.

The samples were added to 200 μl of a 2.5 M solution of hydroxylamine hydrochloride in water, freshly neutralized with NaOH. This mixture was stored on ice until the complete batch could be incubated at room temperature for ten minutes. 0.5 ml of the Fe(III) reagent were then added: 3 % (w/v) FeCl_3_ dissolved in 0.1 M HCl, 12 % (w/v) trichloroacetic acid in water, and 3 M HCl were mixed in equal amounts to obtain this reagent. The reaction was mixed thoroughly and the absorption of the Fe(III) chelates was measured at 540 nm. The absorption values were quantified using a freshly prepared acetyl phosphate standard. Each fraction was measured in triplicates.

#### Statistical evaluation

Statistical evaluations of enzyme activities in the gradient fractions were performed in Origin 8. Based on the assumption of dependent samples, one-way repeated measures ANOVA was performed with an overall significance level of 0.05 and a Tukey test employed for pairwise means comparison.

#### Protein Alignment

Genome search was performed and annotations were taken from JGI IMG/M on 28.6.2019. HMPREF0179_00647 was identified as PduT by Basic Local Alignment Search Tool on UniProt. The search was conducted using the amino acid sequence of HMPREF0179_00647 against UniProt reference proteomes plus Swiss-Prot, with an E-threshold of 10 and no filtering and gapping allowed. For alignments with PduT of *Citrobacter freundii* and *Salmonella typhimurium*, the respective proteins sequences were taken from the UniProt database and aligned with the amino acid sequence of HMPREF0179_00647 using default settings.

## Declarations

### Ethics approval and consent to participate

Not applicable

### Consent for publication

Not applicable

### Availability of data and materials

Not applicable

### Competing interest

The authors declare no competing interest

### Funding

This work was funded by the Deutsche Forschungsgemeinschaft (DFG) (to DS; grant SCHL 1936/4) and the University of Konstanz and Konstanz Research School Chemical Biology (KoRS-CB) (to AGB).

### Authors’ contributions

AGB and DS designed the experiments. AGB, SD, PB and SH performed the experiments and analyzed the data. AGB, PF and CF did the transcriptomic analyses. EI and AK provided support with method development. AGB and DS wrote the manuscript and all authors approved it.

## Acknowledgments

We thank Karin Denger for help with cultivation and Alexander Fiedler, Alina Kindinger and Anne Van Humbeeck for experimental support in Hiwi-projects or during practical classes. We are grateful to Andreas Marquardt for proteomics.

## Notes

### Competing Interest Statement

The authors have declared no competing interest.

