## Supplemental Information for "Bacterial microcompartments for isethionate desulfonation in the taurine-degrading human-gut bacterium *Bilophila wadsworthia*"

**for manuscript:**

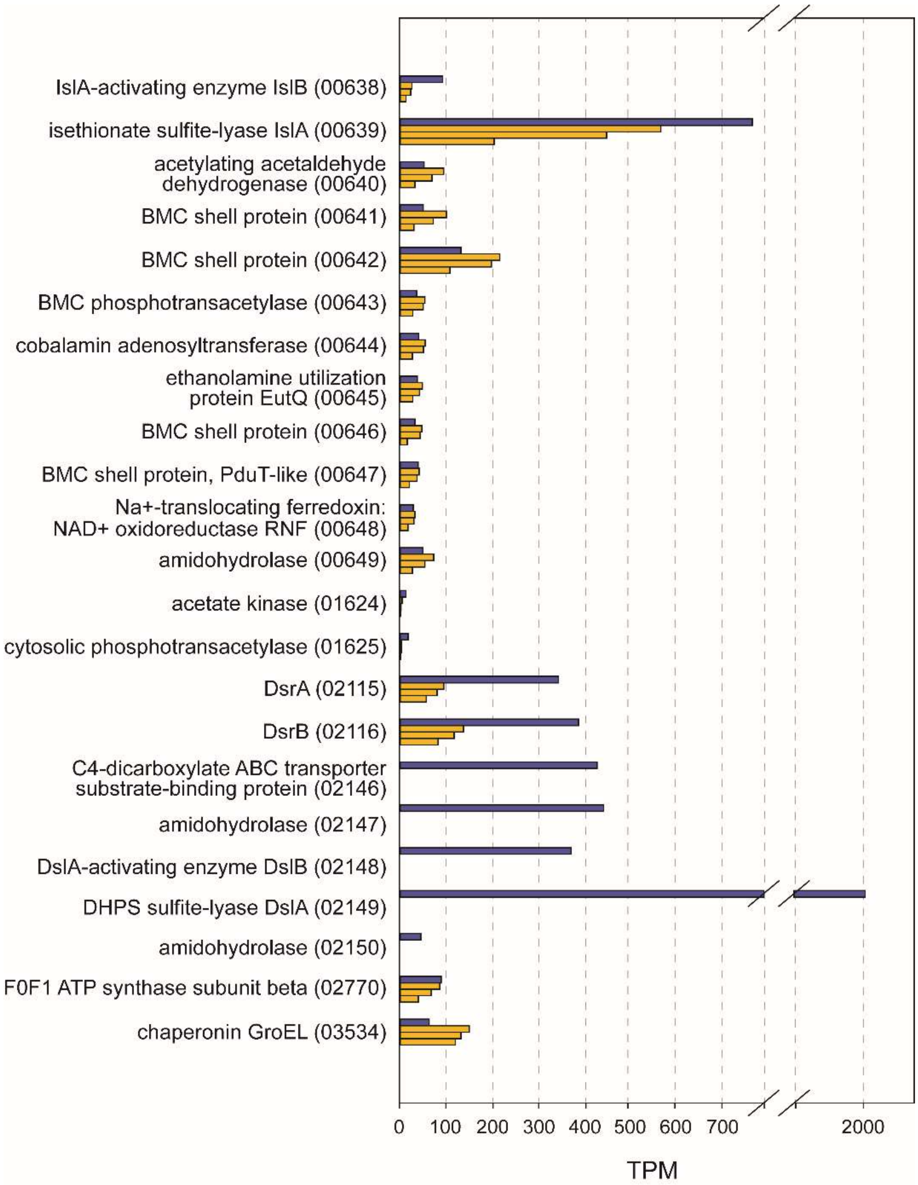


**Figure S1: Transcriptomic data for genes relevant to the discussed pathways.**
Data is shown as transcripts per million reads. Blue bars denote transcriptomic data from a culture grown with 20 mM each of lactate and dihydroxypropanesulfonate, yellow bars from cultures grown with 20 mM each of lactate and taurine (three biological replicates). While taurine is metabolized *via* IslAB (locus tags HMPREF0179_00638 and _00639), DHPS is cleaved by a different enzyme, DslAB (HMPREF0179_02148 and _02149) (i.e., 2,3-dihydroxypropanesulfonate sulfite lyase termed HpsG as most recently described by Liu et al. (1)). Residual expression of taurine-metabolizing genes in the DHPS-grown culture is visible, leading to the assumption that these enzymes are downregulated only slowly when taurine is not present.

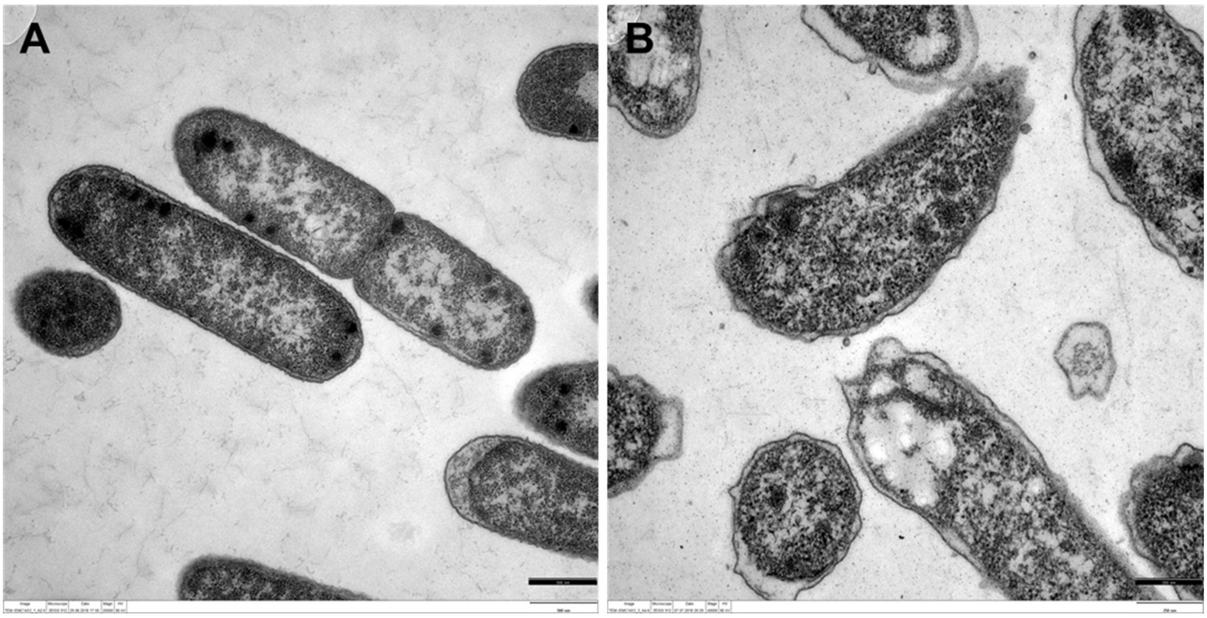


**Figure S2: TEM images of ultrathin sections of *B. wadsworthia* cells grown with taurine (A) or Desulfovibrio alaskensis cells grown with choline (B) as the electron acceptor (B).** Polyhedral structures characteristic of BMCs were observed under both conditions. *D. alaskensis* has previously been described to form microcompartments when degrading choline (2). Scale bars: 500 nm.

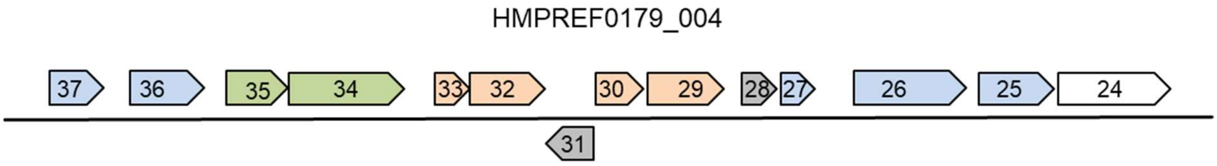


**Figure S3: Illustration of the gene cluster in *B. wadsworthia* encoding sulfolactate degradation enzymes including sulfolactate sulfo-lyase (SuyAB).**
Genes marked in orange are directly involved in sulfolactate degradation: 00433, SuyA; 00432, SuyB; 00430, (S)-sulfolactate dehydrogenase; 00429, (R)-sulfolactate dehydrogenase. The two dehydrogenases most likely interconvert sulfolactate enantiomers, analogous to the pathway described by (3). Genes marked in green encode transport components: 00435, tripartite-type tricarboxylate transporter, receptor component TctC; 00434, putative tricarboxylic transport membrane protein. Genes marked in blue are annotated to encode metabolic enzymes but their function related to sulfolactate metabolism is not clear: 00437, hydroxyethylthiazole kinase (from thiamine metabolism); 00436, Formate dehydrogenase maturation protein FdhE; 00427, methylglyoxal synthase; 00426**,** D-lactate dehydrogenase; 00425**,** glycolate oxidase iron-sulfur subunit. Genes marked in grey may serve a regulatory function: 00431**,** GntR family transcriptional regulator; 00425**,** regulator of RNase E activity RraA. 00424 annotated to encode a hypothetical protein. Notably, there are no genes found in the gene cluster that are annotated to encode BMC shell proteins. All annotations were taken from IMG for *Bilophila wadsworthia 3.1.6*. and the shortened IMG locus tag can be completed by adding the given number to the following prefix: HMPREF0179_.

**Table S1: Number of reads from transcriptomic analysis for RNA samples from *B. wadsworthia* grown with lactate/taurine or lactate/dihydroxypropanesulfonate (DHPS)**

| | Taurine_1 | Taurine_2 | Taurine_3 | DHPS |
|---|---|---|---|---|
| **Raw reads** | 25,780,974 | 21,425,313 | 12,520,332 | 14,901,949 |
| **Filtered reads (% of total raw reads)** | 25,692,021 (99.65%) | 21,347,365 (99.64%) | 12,473,023 (99.62%) | 14,854,536 (99.68%) |
| **reads aligned to reference genome (alignment rate)** | 25,604,277 99.66 % | 21,157,452 99.11 % | 12,430,468 99.66 % | 14,808,153 99.69 % |

**Table S2: Proteomic results for the SDS-PAGE bands (see** Fehler! Verweisquelle konnte nicht gefunden werden.**C). The protein description includes a shortened locus tag, which can be completed by adding the shown number to the following prefix: HMPREF0179_0. BMC shell proteins are marked bold.**

| Fraction | approximate protein size | identified as | Score |
|---|---|---|---|
| **3** | 10 kDa (1) | **ethanolamine utilization protein EutN (0642)** | 115 (top score) |
| | 10 kDa (2) | integration host factor subunit alpha (1532) | 242.03 |
| | | taurine-pyruvate aminotransferase (2713) | 225.79 |
| | | chaperonin GroES (3535) | 196.43 |
| | | DNA-binding protein HU-beta (0718) | 191.42 |
| | | hypothetical protein (0544) | 188.06 |
| | | **ethanolamine utilization protein EutN (0642)** | 157.39 |
| | 25 kDa | dissimilatory sulfite reductase beta subunit (2116) | 655.11 |
| **4** | 14 kDa | LSU ribosomal protein L10P (2084) | 227.30 |
| | 15 kDa | LSU ribosomal protein L10P (2084) | 287.38 |
| | | periplasmic chaperone for outer membrane proteins Skp (2805) | 234.45 |
| | | SSU ribosomal protein S5P (2023) | 234.24 |
| | | LSU ribosomal protein L9P (2745) | 193.54 |
| | | Class III cytochrome C family protein (1235) | 189.18 |
| | | small subunit ribosomal protein S9 (0867) | 187.27 |
| | | SSU ribosomal protein S7P (2067) | 161.72 |
| | | hypothetical protein (2302) | 151.57 |
| | | Flavorubredoxin (3149) | 140.29 |

| | | | |
|---|---|---|---|
| | | **Carboxysome shell and ethanolamine utilization microcompartment protein CcmL/EutN (0647)** | 117.07 |
| | 18 kDa | ATP-dependent Clp protease, protease subunit (2214) | 138.21 |
| | | transcriptional regulator, TetR family (0863) | 135.34 |
| | | hypothetical protein (0984) | 134.65 |
| | | LSU ribosomal protein L9P (2745) | 130.60 |
| | | pyrimidine operon attenuation protein / uracil phosphoribosyltransferase (3139) | 119.28 |
| | | Rubrerythrin (2466) | 95.03 |
| | | Putative exonuclease, RdgC (1649) | 87.47 |
| | | bacterioferritin (0484) | 82.44 |
| | | **Carboxysome shell and ethanolamine utilization microcompartment protein CcmL/EutN (0647)** | 81.44 |

**Table S3: Coomassie and silver staining of SDS polyacrylamide gels, modified after (4, 5).**

| | Substance | time |
|---|---|---|
| **Coomassie staining** | 0.25 % (w/v) Coomassie brilliant blue R-250, 45% (v/v) ethanol , 10% (v/v) acetic acid in water | 1 h |
| **Washing/initial destaining** | 45% (v/v) ethanol , 10% (v/v) acetic acid in water | 1 h |
| **Destaining/Fixation** | 45% (v/v) ethanol , 10% (v/v) acetic acid in water | 14 h |
| **Reduction** | 30% (v/v) ethanol, 0.8 M acetate, 2 g/l thiosulfate | 30 min |
| **Washing** | distilled water | 3x 5 min |
| **Silver solution** | 2 g/l (11.8 mM) silver nitrate, 200 µl/l 37 % formaldehyde | 20 min |
| **Developing** | 25 g/l (0.236 M) sodium carbonate, 100 µl/l 37 % formaldehyde | 2-8 min |
| **Stopping** | 10 g/l glycine | 10 min |

**Table S4: Cell fixation and embedding protocol for transmission electron microscopy**

| | Substance | pH | °C | time |
|---|---|---|---|---|
| **Pre fixation** | 50/50 culture medium with 5% glutardialdehyde in 0.05M HEPES | 7 | 0 | 45 min |
| **Enclosing** | Agarose-enclosure | | | |
| **Fixation** | 2.5% glutardialdehyde in 0,1M HEPES | 7 | 0-4 | 2.5 h |
| **Washing** | 0.05M HEPES | 7 | 0 | 3x 10 min |
| **Postfixation / osmification** | 2% $OsO_4$ in 0.05M HEPES | 7 | 0 | 1h |
| **Washing** | 0.05M HEPES | 7 | 0 | 3x 10 min |
| **Drainage** | 30% ethanol, precooled | | 4 | 10 min |
| | 50% ethanol, precooled | | 4 | 15 min |
| **En-bloc staining** | Uranylacetat saturated in 70% ethanol | | 4 | o.n. |
| **Dehydration** | 70% ethanol, precooled | | RT | 3x 10 min |
| | 80% acetone, precooled | | RT | 3x 10 min |
| | 90% acetone, precooled | | RT | 3x 10 min |
| | 96% acetone, precooled | | RT | 3x 10 min |
| | 100% acetone, dried on molecular sieve | | RT | 3x 10 min |
| **Intermedium** | 100% acetone, dried on molecular sieve | | RT | 1 h |
| **Embedding** | 15% Spurr resin in acetone | | RT | 1 h |
| | 33% Spurr resin in acetone | | RT | 2 h |
| | 50% Spurr resin in acetone | | RT | 2 h |
| | 75% Spurr resin in acetone | | RT | o.n. |
| | pure Spurr resin in closed Eppies | | RT | 2x 2 h |
| **Polymerisation** | | | 65 | 48 h |
| **Cooling** | In extractor hood | | RT | overnight |
